# Essential Dynamics of CB_1_ Receptor-Agonist Complexes: Implications for Signalling Bias

**DOI:** 10.1101/2024.03.25.586709

**Authors:** Daniel M. J. Fellner, Michelle Glass, Daniel P. Furkert

## Abstract

The CB_1_ cannabinoid receptor is implicated in a broad range of physiological processes and disease states, however CB_1_-targeting drugs in clinical use remain based on tetrahydrocannabinol (THC). Ligands that exhibit functional selectivity for different intracellular signalling pathways are currently an area of rapid development, and hold significant potential as therapeutic agents. Improved understanding of the exact molecular mechanisms underpinning biased activation of intracellular effector proteins for CB_1_ is necessary to enable drug development of biased CB_1_ ligands into candidates for treatment of human disease. Using molecular dynamics, this study shows that CB_1_conformations resulting from activation by the orthosteric ligands CP55940, Δ9-THC or 5F-MDMB-PICA exhibit differences in the dynamic organisation of key residues and receptor substructures involved in coupling to G proteins and β-arrestins, that leads to selective activation of downstream signalling pathways. The identification of conformationally distinct CB_1_-agonist complexes that demonstrate different functional profiles provides an important step in unravelling the molecular determinants for biased signalling, and lays a platform for future rational design of novel therapeutic leads.

## Introduction

The cannabinoid CB_1_ G protein-coupled receptor (GPCR) is primarily expressed in the central nervous system, and as part of the endocannabinoid system (ECS) is linked with a number of metabolic or neurodegenerative processes and disorders including obesity, depression, learning, pain, Parkinson’s disease and cancer^.1,2^ Many studies have explored the potential therapeutic applications of CB_1_ ligands,^3^ but these efforts have been hampered by the incidence of adverse effects including psychiatric disorders.^4^ The CB_1_ receptor has been understood to act though the canonical Gα_i_-mediated signalling pathway and β-arrestins, however signalling via other G proteins has also been demonstrated.^5^ Importantly, it has been established that ligands for CB_1_ activate intracellular signalling pathways to a different extent, a concept termed ‘biased signalling’ or ‘functional selectivity’^.6,7,8^ Evaluation of unique signalling profiles elicited by established CB_1_ ligands across different signalling pathways has recently generated strong research interest.^9,10,11,12^ Kinetic modelling of CB_1_ signalling bias,^13^ and signalling profiles for recently reported classes of synthetic cannabinoids have also been reported.^14,15^ Increased fundamental understanding of the molecular determinants driving ligand signalling bias is important to enable development of drug candidates that modulate specific processes via CB_1_with minimal undesired activity.^16,17^ Despite the availability of increasingly detailed structural information through crystallography and cryo-EM, the dynamic mechanisms underpinning functional selectivity in cannabinoid receptors remain poorly understood. Molecular dynamics (MD) simulations provide a complementary approach for analysing receptor function and are increasingly important in identifying signal transduction events.^18^ In this study, MD was used to study the conformational dynamics of the CB_1_ cannabinoid receptor complexes of three agonist ligands, to explore possible mechanisms for biased signalling towards G proteins and β−arrestins. Mapping of receptor ligands to quantifiable structural differences relies on consistent and comparable signalling data. At the outset of this study, this had been provided by a recent comprehensive analysis of CB_1_agonist functional selectivity reported by the Glass group.^19^ To capitalise on the availability of this comprehensive data set from a single source, three of the ligands evaluated were chosen for our computational investigations, to span a range of observed signalling bias (**Figure 1**); Δ^9^-THC, a partial agonist which signals via G proteins, but had undetectable signally via β-arrestins; CP55940, an agonist commonly used in pharmacological research with a relatively balanced signalling profile; and 5F-MDMB-PICA, a synthetic cannabinoid receptor agonist (SCRA) more biased toward G protein signalling than CP55940.

**Figure 1.**
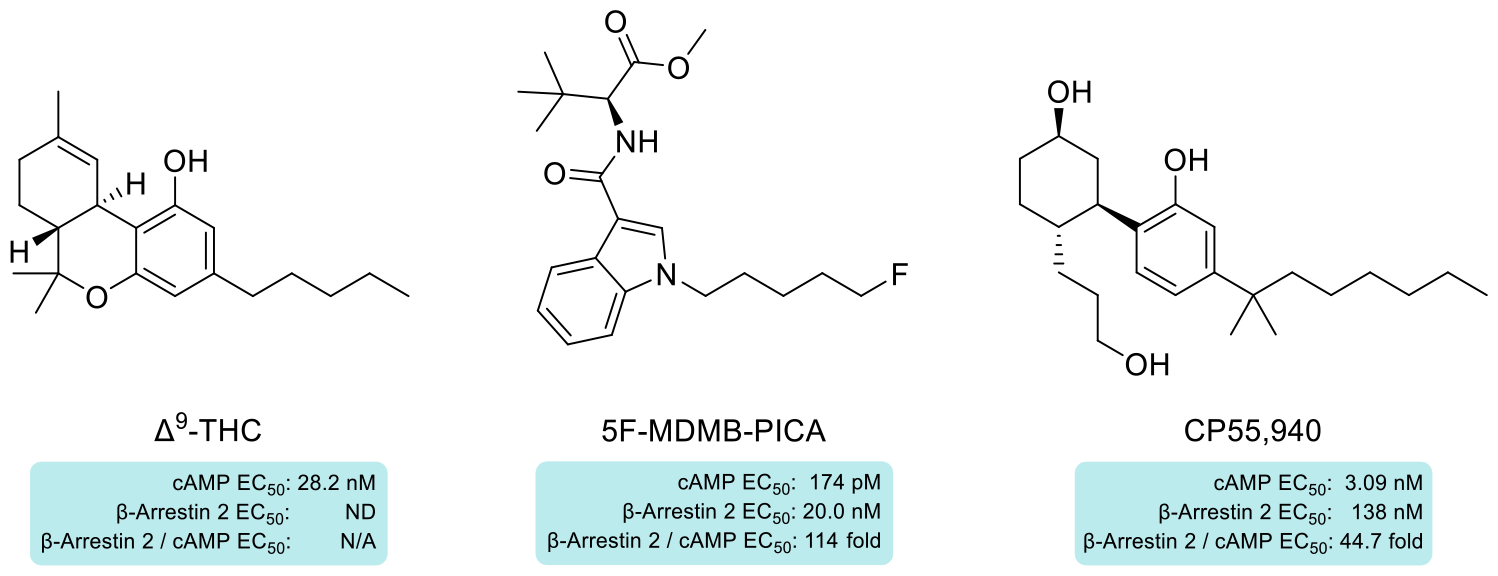
Orthosteric ligands of CB1 with well-established diverse signalling profiles investigated in this study.

In the current study, molecular dynamics (MD) simulations of the receptor were used to investigate whether the binding of different ligands led to significant differences in receptor conformational dynamics, in a preliminary investigation towards understanding the molecular basis for biased signalling of orthosteric CB_1_ agonists (**Figure 2**). MD simulations are well established to provide important complementary *dynamic* information to that obtained through crystallography and cryo-EM structures that reveal precise, high-resolution structural detail, from single *static* images.^20,21^

**Figure 2.**
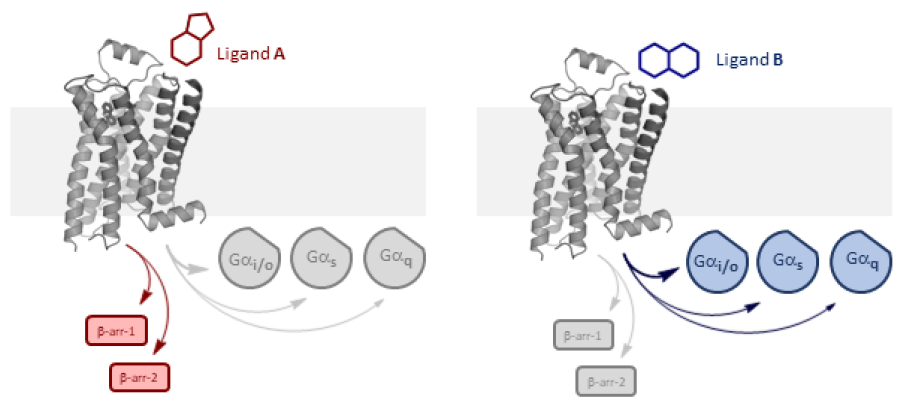
Simplified biased ligand signalling at CB_1_ for β-arrestin vs G protein pathways.

## Methods

A model of hCB_1_ was assembled based on an active state cryo-EM structure,^22^ guided by literature knowledge of active state conformational determinants.^23^ This receptor was initially equilibrated in the apo form in a heterogeneous bilayer for 1 μs. The three test ligands were docked, ligand force field parameters were optimised and validated, and each ligand-bound CB_1_complex was simulated for 600 ns, in duplicate (3 ligands, 2 runs for each, of 600 ns duration). The first 100 ns of each simulation were discarded to allow for equilibration, and the trajectories were aligned for principal component analysis (*see* Supplementary Data for details).

## Results and Discussion

### Characteristic occupancy of the orthosteric binding pocket for CB_1_-agonist complexes

Quantification of the average spatial density^24^ for each of the test ligands in the CB1 orthosteric binding pocket, from the same starting geometry, showed that the orthosteric binding site adopted distinctly different conformations in binding to each of the test ligands, with some expected variation between the duplicate pairs of simulations (**Figure 3**).

**Figure 3.**
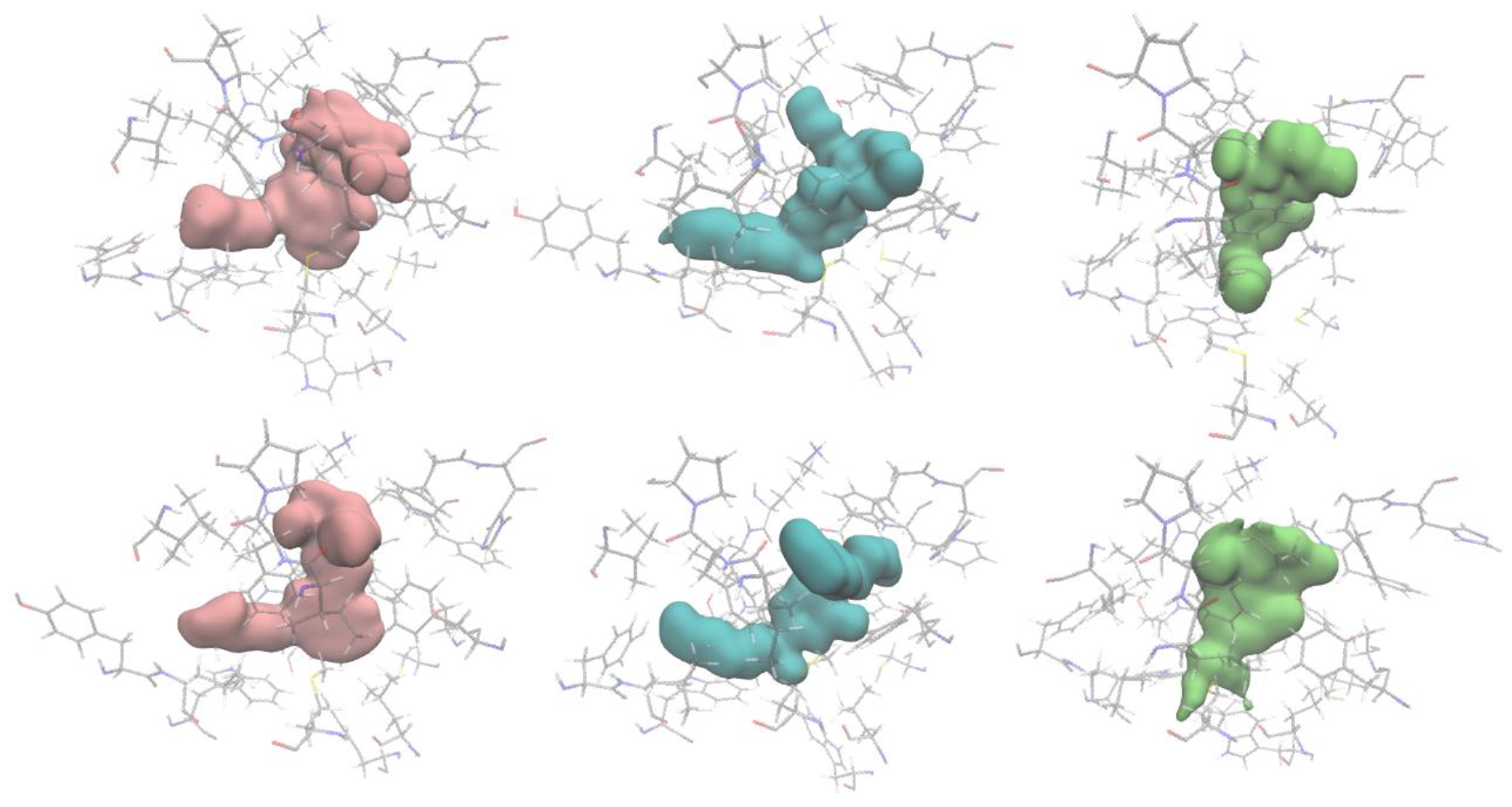
Average density maps (one image for each duplicate simulation of the three ligands) for 5F-MDMB-PICA (5MP), CP55,940 (CP) and Δ9-THC (THC) in the CB_1_ orthosteric binding site.

### Principal component analysis (PCA). General introduction

The spatial differences in the simulations observed spanned many amino acid residues, widely distributed across the transmembrane domains of the receptor. To usefully interrogate this multidimensional data, principal component analysis (PCA) was employed.^25^ PCA is a tool applied in the analysis of complex systems, where the large number of variables limits ready extraction of useful information. This technique has previously been applied to the study of GPCR dynamics^.26,27^ In simple terms, the approach involves identification of virtual vectors through the dataset, called principal components, whose variation is responsible for a portion of the variability observed for the whole system. There are many such principal components, assigned as PC1, PC2, PC3 *etc*, in order of decreasing importance. In practice, only the first few principle components are usually considered (reference!). This analysis is able to characterise features or differences of *datasets*, and can be analysed to determine the main conformational contributors to the principal components. For the present study, graphical analysis of the principal components of the simulation data, namely the receptor’s cartesian coordinates, enabled the identification of similarities and differences in the conformational dynamics of the CB_1_-ligand complexes, down to the level of receptor sub-domains or individual residues.^28^ Each data point in these graphs represents a single frame of the simulation, placed according to its coordinates on the named principal components. The combined data points for all frames over the simulation thus allow graphical representation of the dynamic variation of the system under study.

### PCA of data for CB_1_-ligand complexes

Once regions exhibiting significant differences between the three CB_1_-ligand complexes had been indicated by PCA, it was possible to extract and analyse relevant *physical* data from the atomic coordinates in the frames of the simulations (*e*.*g*. bond angles, interatomic distances, helix conformation, ligand-receptor contacts) to build an understanding of the differences in dynamic behaviour between the three receptor-agonist complexes.

### PCA of orthosteric binding site residues

Initially, analysis focused on the residues of the CB_1_ receptor making up the orthosteric binding site.^29^ Graphical inspection of principal component data showed significant differences between the three test ligands (**Figure 4**). Points corresponding to the two full agonists (5MP and CP5) were clustered close together in the PC1,2 plot (*left*) while partial agonist (THC) gave two separate clusters, expected to correspond to active and inactive CB_1_ conformations. This suggests that the PC1,2 plot likely describes binding site dynamics governing differences in canonical receptor activation. In contrast, the distribution of datapoints into the more disperse clusters observed in PC3,4 for 5MP, CP5 and THC (**Figure 4**, *right*) suggested that these represented more subtle conformational variations in receptor dynamics, mediating more differential transducer binding.

**Figure 4.**
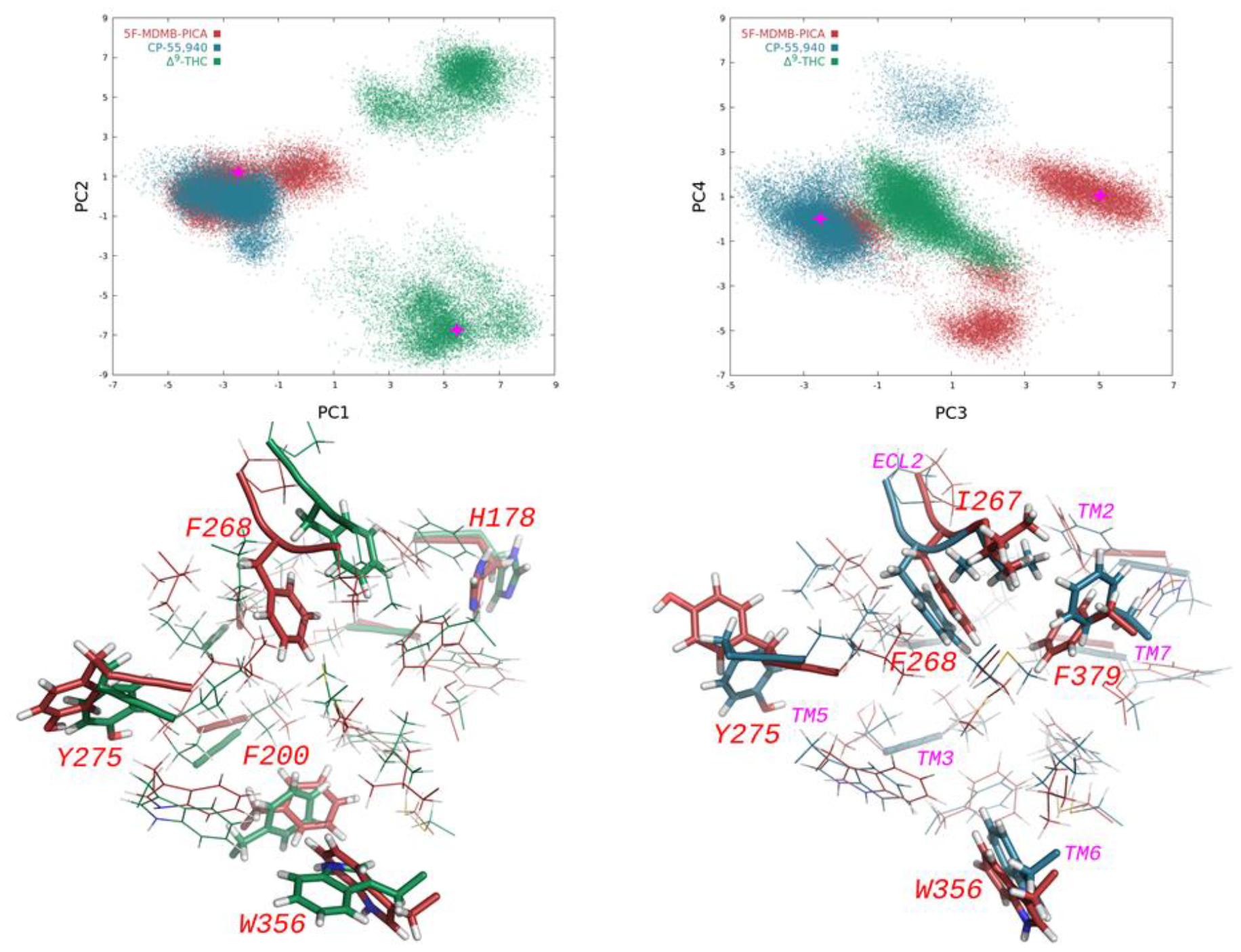
Lateral view of the orthosteric binding site with TM5,6 in the foreground. Overlay of simulation frames from PC1,2 for CB1-5F-MDMB-PICA vs CB1-Δ9-THC (*lower left*), and CB1-5F-MDMB-PICA vs CB1-CP55,940 from PC3,4 (*lower right*), showing major modes of variation, with key residues highlighted. PCA plots (*top left* and *top right*, note pink points) indicate the source of the respective simulation frames shown in the respective overlays below.

### Ligand interactions with specific residues in the receptor orthosteric binding site

In order to probe interactions at the binding site more closely, the percentage of simulation frames showing contact (<4Å) between the ligand and residues in the binding site was quantified.

Data from individual frames of each simulation was used to establish a percentage time that specific residues were located within 4Å of the respective test ligands. Data for a selection of notable amino acids is shown in **Table 1**, and is shown grouped by sub-domain in **Figure 5**.

**Table 1.** Residue contact % at the CB_1_ orthosteric binding site (ligand within 4Å).

|  |  | Δ9-THC | 5MP | CP55,940 |
| --- | --- | --- | --- | --- |
| TM2 | F170 | 92 | 99 | 100 |
|  | F174 | 49 | 47 | 100 |
|  | H178 | 15 | 29 | 96 |
| TM3 | F189 | 81 | 89 | 73 |
|  | K192 | 80 | 71 | 50 |
| EC2 | I267 | 3 | 92 | 67 |
|  | F268 | 100 | 98 | 99 |
|  | I271 | 39 | 81 | 97 |
| TM5 | Y275 | 7 | 83 | 99 |
|  | W279 | 98 | 100 | 100 |
| TM6 | L359 | 100 | 99 | 92 |
|  | L360 | 66 | 3 | 48 |
|  | M363 | 98 | 100 | 99 |
| TM7 | F379 | 100 | 100 | 99 |
|  | A380 | 1 | 1 | 42 |
|  | C386 | 60 | 73 | 6 |

**Figure 5.**
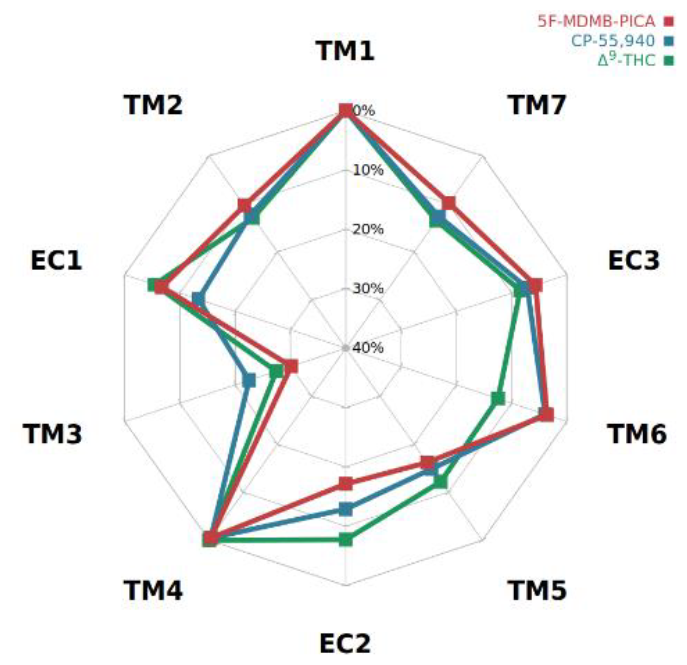
Ligand-receptor interaction energy grouped by sub-domain as percentage of ligand total.

Notably, the ligands were found to differ substantially in their fractional contact time with residues on TM2, TM7, and EC2. Analysis of ligand interaction energies with the binding site sub-domains also revealed considerable variation in the relative contributions of residues from TM3, TM6, and EC1 to their binding. Each ligand displays a unique interaction profile with the orthosteric binding site residues, providing a quantifiable basis for ligand-specific stabilisation of distinct receptor conformations driven by structural changes in the binding site.

### Conformational dynamics for receptor subdomains

More specific PCA was next used to probe the dynamic variability of the individual receptor sub-domains, for the three CB_1_-ligand complexes. Graphical representation of PCA data (**Figure 6**) showed that transmembrane helices TMH2, TMH3, TMH5 and TMH7 generated the most widely-spread PCA clusters. This suggested that these varied the most between the CB_1_-ligand complexes, and accordingly might contribute significantly to the conformational changes responsible for the molecular mechanism of agonist bias. Based on the crystallographic and cryo-EM structures of CB_1_ and other GPCRs bound to G proteins or β-arrestins, residues from TMH 2, 3, 5, 6 & 7 and the intracellular domains ICL2 and ICH8 form key interactions with these transducer proteins.^30^

**Figure 6.**
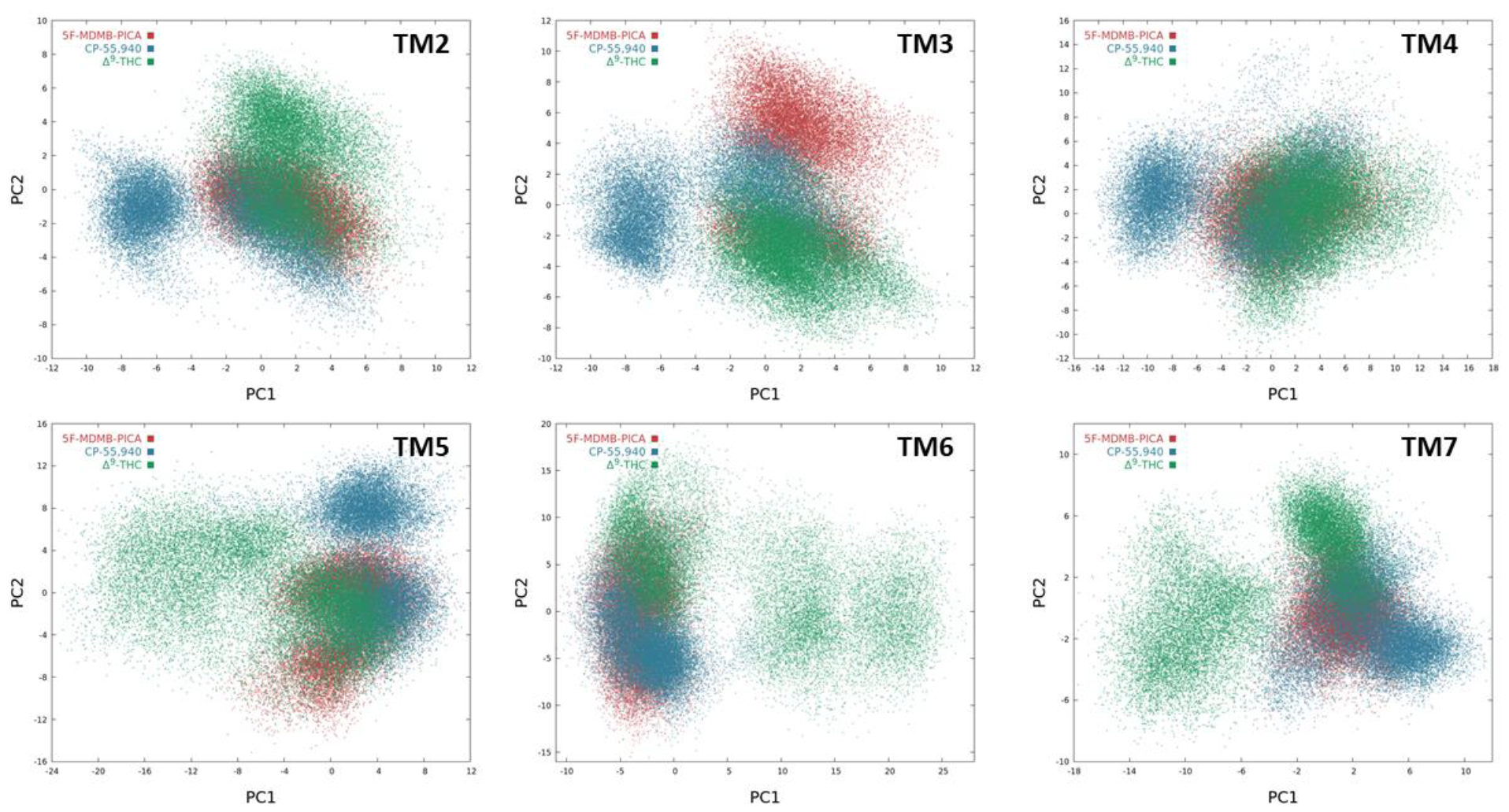
Principal component analysis revealed distinctive differences in the dynamics of the individual transmembrane helices, between each of the three CB1-agonist complexes; 5F-MDMB-PICA, CP55,940 and Δ9-THC (Data for each domain is fitted to the entire receptor).

The existence of multiple, sometimes relatively diffuse clusters of the ligand THC on the PC projections suggests that THC stabilises a relatively broader range of conformations in these sub-domains. In conjunction with the fact that these diffuse clusters are well-separated from clusters of the full agonists 5MP and CP5, these THC clusters likely represent inactive or less-active receptor conformations. Encouragingly, the full agonists 5MP and CP5, which differ somewhat in their signalling profiles with respect to G protein or β-arrestin2 recruitment, are also differentiated along the principal components; particularly in subdomains TM2, TM3, TM5, and TM7. The signalling profile of CP5 is generally considered by pharmacologists to be well-balanced, in contrast to 5MP which was selected for this study due to its combination of high efficacy and relative bias for cAMP inhibition over β-arrestin2 recruitment (ref). On the PC projections of subdomains TM2, TM4, and TM7, 5MP appears predominantly as a single cluster, whereas CP5 explores two or more clusters in these sub-domains. A possible explanation is that the more numerous clusters in these sub-domains are reflective of the relative signalling diversity of CP5.

These data reflect dynamics of the transmembrane helices relative to the receptor corresponding to larger scale motions such as lateral or vertical displacements with respect to the bundle of helices. In order to probe smaller-scale movements within the helices, for each of the receptor sub-domains analysed in **Figure 6**, further PCA was carried out based on the least-squares fit of each domain to itself, rather than the whole receptor. Selection of two representative physical conformations, corresponding to opposite ends of the principal components, enabled identification of residues whose movement contributed significantly to those principal components, and the nature of the motions made by each one (**Figure 6**).^31^ Quantification of relevant data for these key residues of interest, *e*.*g*. F379 in TM7, and Y153 in TM2 (*see* **Figure 9** histograms) allowed additional comparisons to be made between the three agonist-CB_1_ complexes.

### Residues contributing most to PC1 and PC2 for each transmembrane domain

Within the receptor sub-domains, the residues that contributed most to the principal components 1-4 for each receptor subdomain were identified (**Table 2**).

**Table 2.** Residues contributing most to PCs 1-4 of the TM domains, in decreasing order of importance (*see* SI for further details).

|  |  |
| --- | --- |
| TM1 | Q115, L138, Q116<br>F129, L126, L136, I119, L117 |
| TM2 | **F170, Y153, F174, Y172**<br>I169, I175, I165 |
| TM3 | **F191, N187, F189, K192**<br>R214, Y215, V188 |
| TM4 | **F237, R230, I245, I247, L239, M240, V249, W241, K232** |
| TM5 | H302, H304, V306, **Y275**<br>F289, L276, F278, W279 |
| TM6 | R336, L359, L360, R340, **W356**, M337 |
| TM7 | **Y397, F379, F381**<br>I396, L385, L388, V392, N393 |
| Important residue in PAM exosite proposed by Hurst *et al.*<sup>31</sup><br>† Receptor activation toggle switch F200-W356<br>Homologous residues implicated in GPCR agonist bias<br>Bold residues appear in main text analysis<br>Grey residues appear <3 times |  |

Among this group of amino acids, several (*in purple*) were homologous to residues previously implicated in biased signalling for other GPCRs; Y275, F278, W279, M363, F379, L387 and Y397.^32,33,34^ Of particular interest, aromaticity at position Y275^5.39^ has been established as critical to cannabinoid ligand binding by mutation studies; CB_1_ mutant Y275F displayed similar pharmacology to the wild type, however the mutant Y275I did not bind CP55,940 due to rearrangement of the orthosteric ligand binding pocket, with loss of signal transduction.^35^ Aromatic residues occupy the homologous position in related GPCRs (CB2 T190^5.38x39^, βAR Y172^5.38x39^, β2R Y199^5.38x39^, A2A Y211^5.38x39^, D2 F189^5.38x39^, D4 Y192^5.38x39^) and evidence suggests these play a key role in ligand recognition and biased signalling (references). The D2 mutant F189Y has demonstrated increased affinity and altered subtype selectivity for specific ligands.^36^ More crucially for the present study, structural permutations in the mutants D2 F189A and β2R Y199A were shown to propagate allosterically to the intracellular surface, affecting interactions with transducers to mediate signalling bias.^37^ Tyrosine Y172^2.59^ in TM2 is also of interest, potentially playing an important role in the allosteric binding site of the ago-PAM GAT229 proposed by Hurst et al._38_

Visualisation of the conformational changes described by the main principal components by viewing receptor structures at different coordinates along the PCs revealed several major changes in both extra- and intracellular residues across a number of sub-domains (**Figure 7 & 8**). This allowed the evaluation both of differential agonist stabilisation of the binding site, and the resulting allosterically transmitted effects in the intracellular transducer-binding cavity.

**Figure 7.**
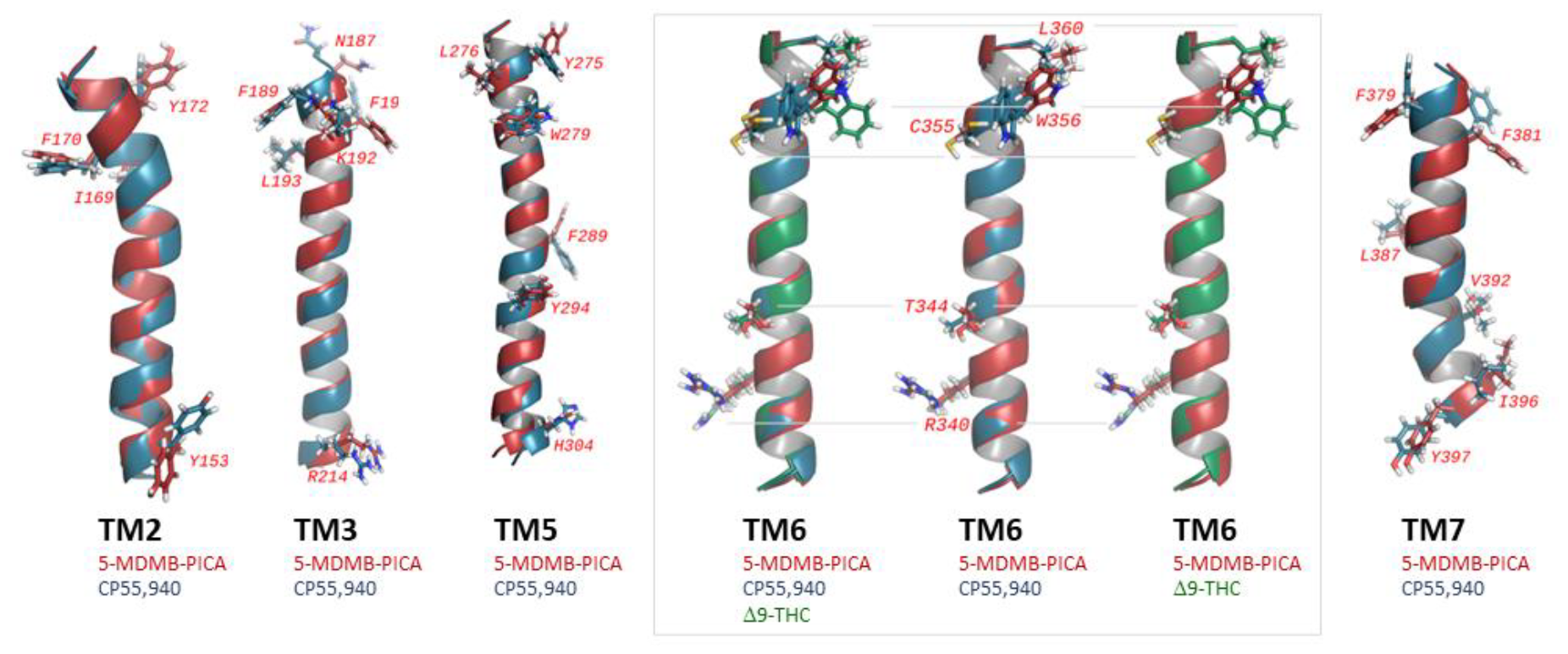
Overlaid pairs of transmembrane helix conformations, showing differences in residue orientations for the complexes CB1-CP55,940, CB1-5F-MDMB-PICA or CB1-Δ9-THC, at opposite ends of the noted principal components; TM2 CP-5MP (*far left*, PC2, note Y153); TM3 CP-5MP (*second from left*, PC1, note F191, N187, R214); TM5 CP-5MP (*third from left*, PC2, note Y275, F289); TM6 (*grey box*, CP-5MP-THC, CP-5MP, 5MP-THC; note R340), TM7 CP-5MP (*right*, PC2, note F379, F381).

**Figure 8.**
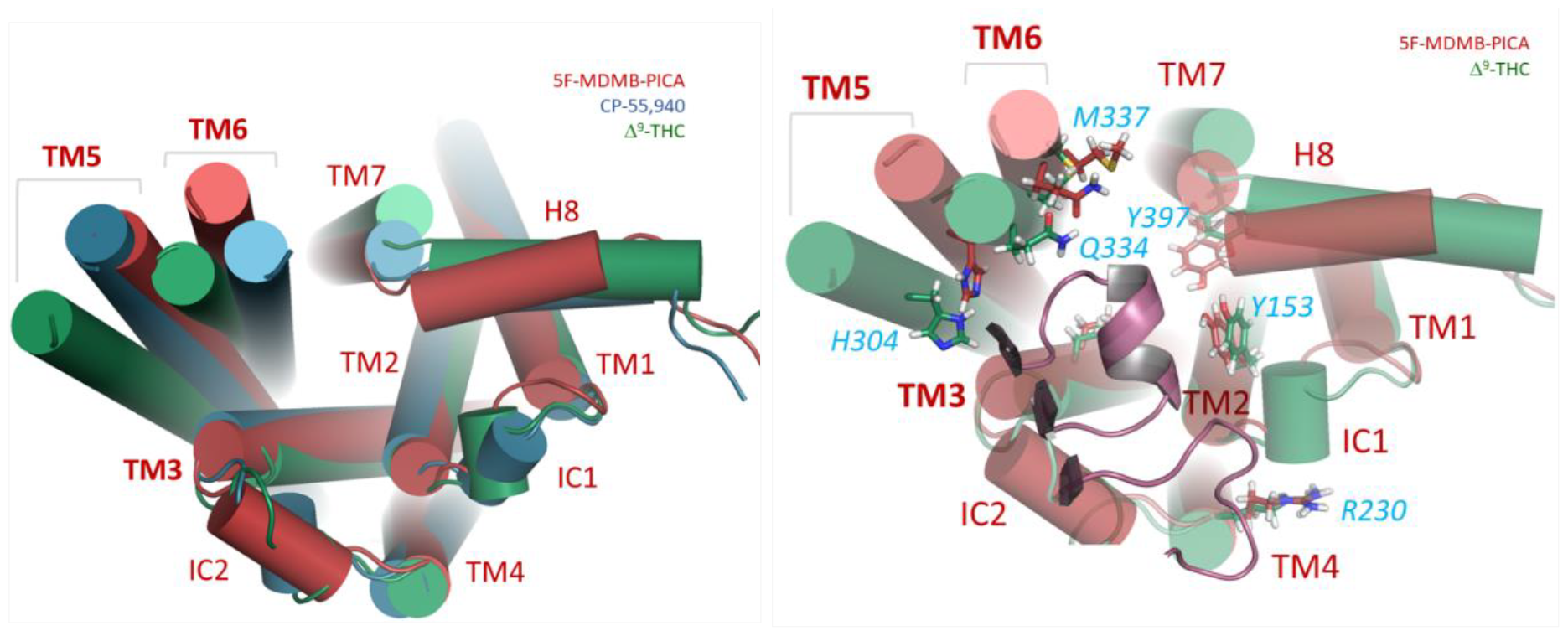
(*Left*) Conformational differences at the intracellular receptor surface containing the transducer binding pocket, for structures representing opposite ends of PC2 (for the entire transmembrane domain of the receptor) for all three CB1-agonist complexes; (*right*) CB1-5F-MDMB-PICA and CB1-Δ9-THC complexes overlaid, with significant residues highlighted.

The entire transmembrane domain was also analysed by PCA, to determine and visualise the conformations stabilised by the ligands in the context of all the helices (**Figure 9**, *left*). This revealed that on the intracellular side, displacement of TM5 and TM6 were the largest conformational changes seen between the agonist-receptor complexes. Although the simplified representation in Figure 7 omits subtleties in the conformation of TM7 between the full agonists 5MP and CP5, they produce different amounts of lateral displacement in H8 and the residues connecting TM7 to H8. This region is hypothesised to be important in the binding of β-arrestin-2 to CB_1_. Lateral displacement of this element of the receptor alters the intracellular cavity surface available for transducer binding, in this region.

**Figure 9.**
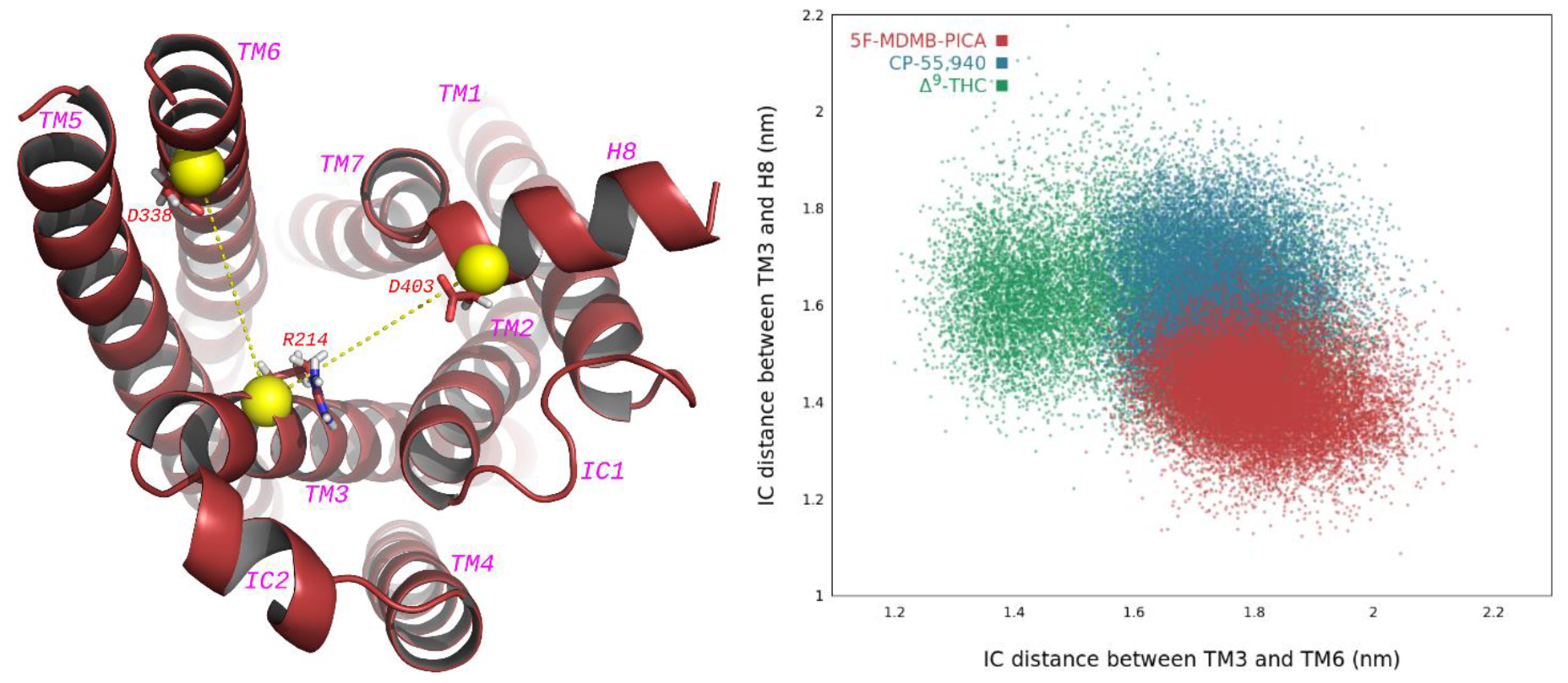
TM3-TM6 vs TM3-TM7/H8 distances at the intracellular surface (*defined at left*) showed characteristic clusters for the three ligand-receptor complexes (*right*).

IC1 and IC2 were also shown to differ substantially in conformation between the principal components. Notably, experimental structures of the M2 muscarinic acetylcholine^39^ and rhodopsin^40^ receptors bound to either Gα_o_ or β-arrestin1 show conformational differences in these regions (as well as in TM5, TM6 and the TM7/Helix 8 hinge region), and that IC2 forms direct interactions with both Gα_o_ and β-arrestin1.

### Distances from TM3 for characterisation of the transducer binding pocket

To analyse the spatiotemporal distribution of the main motions contributing to transducer binding pocket dynamics, the relatively stable helix TM3 was used as a reference to measure the extent of pocket opening at TM6 and the TM7/H8 interface. Plotting these in two dimensions (**Figure 9**, *right*) allowed estimation of the relative shape of the cavity stabilised by the ligands. Differences in receptor activation between THC and the full agonists are reflected in the distances between TM3 and TM6, while the opening of the cavity near the TM7/H8 interface differs between the full agonists.

### Comparison of conformational data for selected residues around the transducer binding pocket

With the most variable residues identified, more granular analysis such as rotamer preferences was used, to examine how spatiotemporal arrangement of residues differed between ligands. For example, F379^7.35^ may interact with M363^6.55^ and ECL2 in one rotamer, or F177^2.64^ and H178^2.65^ in another. This shift may influence helix positioning, particularly between TM2 and TM7. Both helices have been implicated in the binding of transducer proteins. The rotamer of F379^7.35^ favoured by 5MP is adopted at the same time as a unique rotamer of L387^7.43^, that sits wedged between TMs 1, 2 and 7. The latter residue has also been implicated in agonist bias.^41^ The dihedral angle defining the Y397 rotamer of the N^7.49^P^7.50^xxY^7.53^ motif (**Figure 10**, *left*) was observed largely in a single state, in line with the recently reported CB1-βarrestin1 complex that noted the role of Y397, and the T210^3.46^, Y294^5.58^, Y397^7.53^ triad polar network in recruiting βarr1.^42^

**Figure 10.**
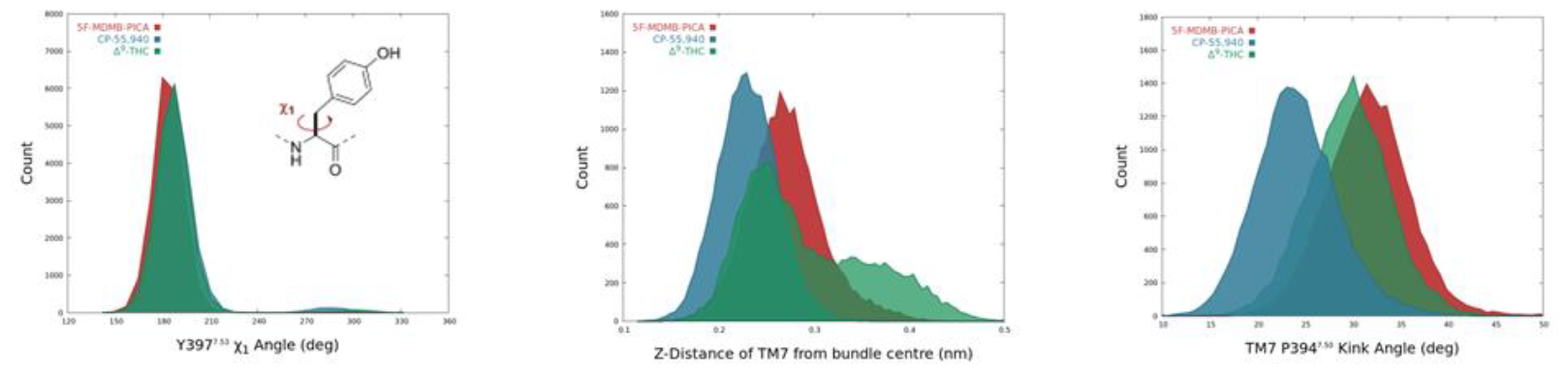
(*Left*) Comparison of Y397 dihedral χ_1_ in the three CB_1_-agonist complexes. Differences in TM7 dynamics influential for the transducer binding pocket were observed, notably the TM7 vertical position (*centre*), and particularly for the TMY kink angle at proline P394 (*right*).

that emphasized the key role of Y397 rotamers, and the dependent T210^3.46^, Y294^5.58^, Y397^7.53^ triad polar network in recruiting βarr1.^43^ The NPxxY motif generally, including proline P394^7.50^, is heavily implicated here as a significant contributor to the molecular mechanism of bias transmission, by data in this study. Tyrosine Y397^7.53^ is one of the major contributors to the TM7 principal components. The TM7 kink that is observed to be straightening in the CB_1_-CP55940 complex (**Figure 11**) is bending at P394^7.50^. Divergence in the D163^2.50^ to N393^7.49^ H-bonding status was also observed between the three CB_1_-ligand complexes.

**Figure 11.**
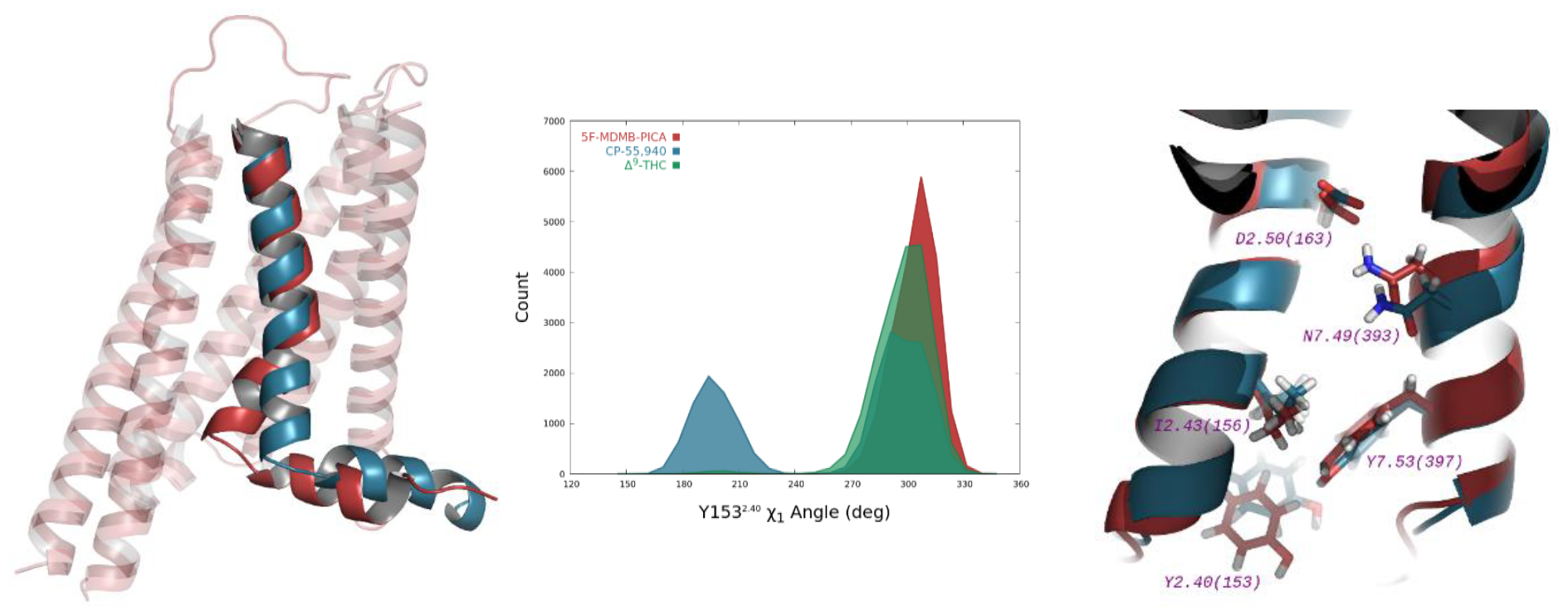
Detailed views of variations in TM7 position between CB_1_-5F-MDMB-PICA and CB_1_-CP55940. Of particular note are the TM7 kink angle and its influence on defining the size of the transducer binding pocket (*left*, red vs blue), and the consequences of Y153 conformation (*right*).

### W279-M363 Distance

It has been reported that for Family A GPCRs, residues 6.55 and 5.43 are functional hotspot residues that contribute significantly to allosteric communication pipeplines linking the agonist binding site to the transducer interacting interfaces,^34^ but the homologous residues in CB_1_ (Met and Trp) are incapable of directly analogous H-bonding. In some clusters of the 5MP trajectory in the current study though, they do get close enough to interact (**Figure 12**).

**Figure 12.**
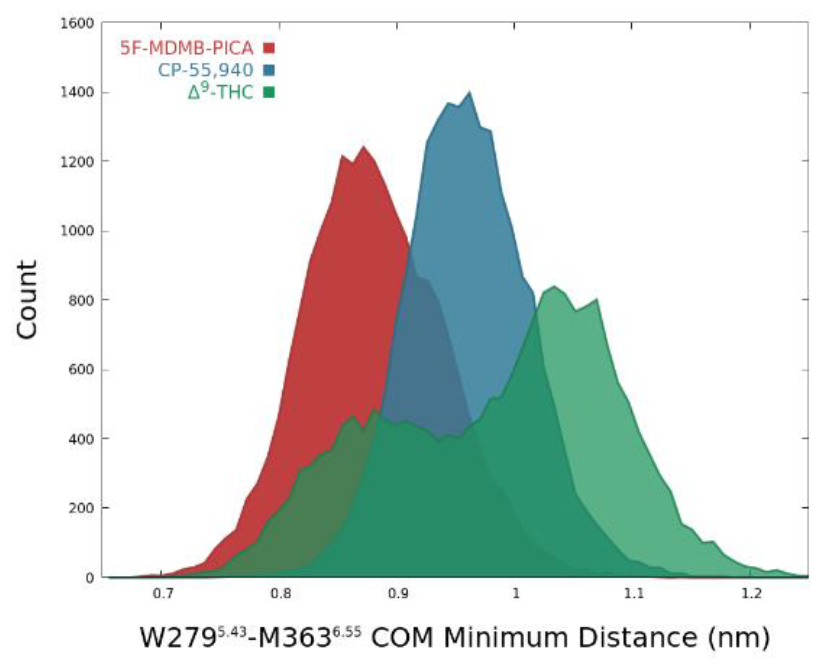
Comparison of W279-M363 distance in CB1-ligand complexes.

### Pentane tail orientation

The pentane tail in the CB_1_-Δ9-THC complex was not observed to extend as far into the hydrophobic cavity, as for the CB_1_ complexes with 5F-MDMB-PICA or CP55940 (**Figure 13**). THC only interacted transiently with the tail pocket residues (*e*.*g*. W279^5.43^ and Y275^5.39^), in contrast to the full agonists. This may explain the observation that THC-bound CB_1_ recruits minimal β-arrestins, as our data implicates these residues in agonist bias. The homologous residues in other receptors have previously been found to govern agonist bias as well. If this holds for other partial agonists, it may explain their relative lack of β-arrestin recruitment too. These residues, including Y275^5.39^ in particular, also show different rotamer profiles between the full agonists, possibly reflecting their differing β-arrestin bias.

**Figure 13.**
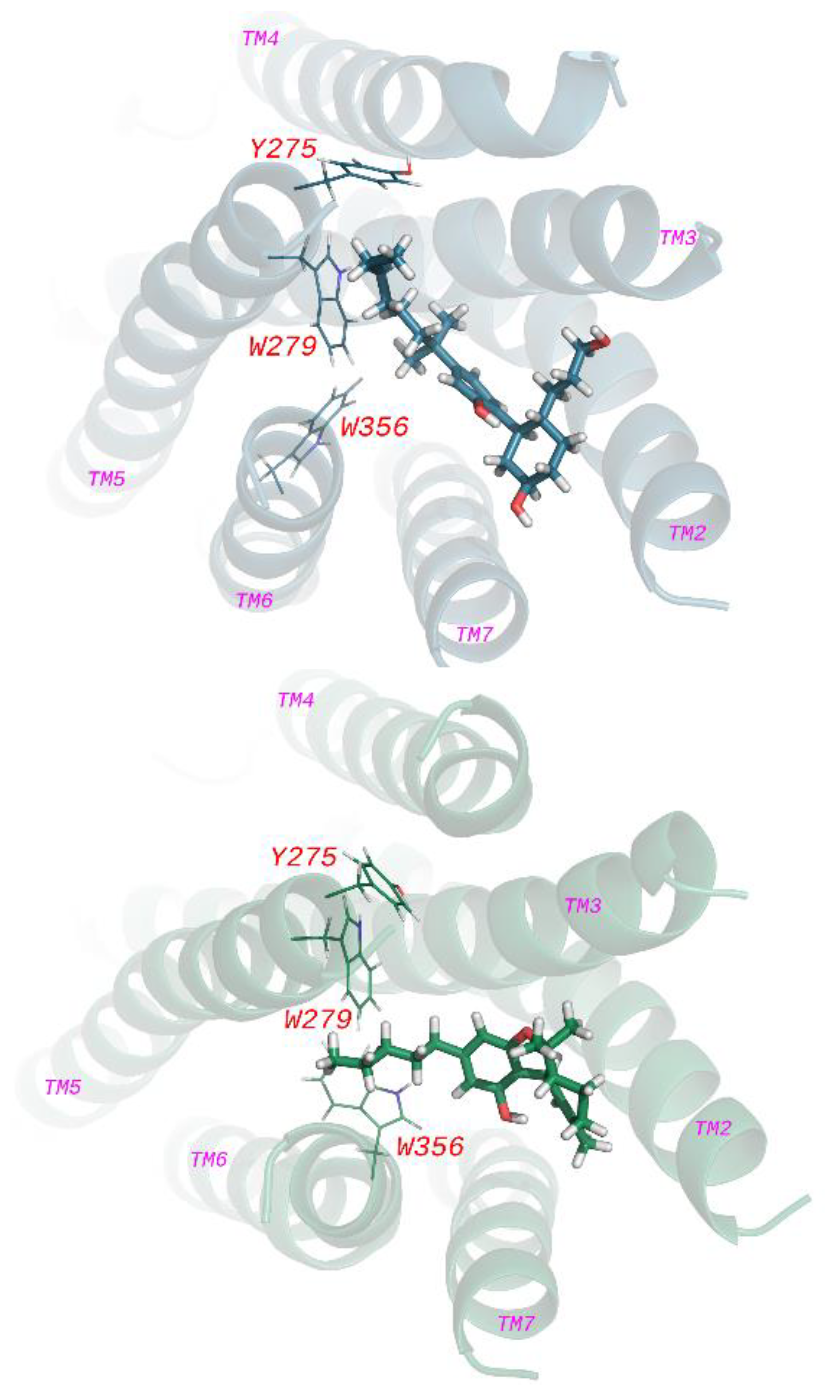
Comparison of ***pentane tail orientation*** between the CB_1_-CP55940 complex (*top*) and CB1-Δ9-THC (*bottom*), highlighting the differing positions of key residues W279^5.43^ and Y275^5.39^.

It should be noted that the relative *potency* of CP5 for β-arrestin vs G proteins is higher than for 5MP, but its *efficacy* at recruiting β-arrestin is far lower (−50%) than that of 5MP. During simulations, the Y275^5.39^ residue was observed to alternate between two rotamers, giving a distinct distribution histogram for each ligand.

## Discussion

Potentially, signalling bias may be a result of the relative time spent in each cluster by the respective ligands, as this could correlate to differences in transducer affinity for the respective conformations. The other case is that biased signalling conformations are geometrically unique for each ligand, with each of these being a constrained conformational subset of the active conformations of the respective receptor-ligand complex. Future simulations of the receptor bound to G proteins or β-arrestins, may enable further deconvolution of PCA clusters, to identify the contributions of these geometrical or temporal effects to the observed signalling bias. Additionally, given that ligand and transducer binding are dynamic equilibrium processes, it is possible that transducer binding pre-disposes the receptor for biased signalling, by inducing conformational changes that favour binding and activation by a particular orthosteric ligand.^44^ Alternatively, some part of the measured signalling bias may be due to unique differences in receptor number on the cell surface, expression of the effector proteins, or the temporal profile of ligand binding and the elicited signalling.^45^

While a range of GPCR oligomerisation interfaces have been proposed, the involvement of TM5 is a recurring theme in the formation of GPCR homo- and hetero-oligomers.^46,47^ Additionally, synthetic CB_1_ TM5 and TM6 peptides were reported to disrupt receptor heteromerization with 5-HT_2A_ receptors.^48^ The same study reported that 5-HT_2A_-knockout mice did not develop cognitive impairment when undergoing treatment with THC.^49^ Differences in the conformational dynamics of TM5 in the presence of different ligands, *as seen in this work*, may therefore influence the extent of receptor oligomerisation and/or the type of oligomers formed by CB_1_, leading to different signalling outcomes.

## Conclusions

Molecular dynamics simulations of the CB_1_-ligand complexes of 5F-MDMB-PICA, CP55940, and Δ9-THC revealed characteristic conformational dynamics profiles for each ligand. Principal component analysis was used to identify specific sub-domains and residues of the receptor likely to contribute to biased signalling via G proteins or β-arrestins. Detailed comparisons of ligand-receptor contacts, amino acid conformer preference and transmembrane helix conformation were able to be made between the three ligand-receptor complexes. Several residues highlighted in this study were homologous to those already implicated in determination of bias for other GPCRs. Molecular dynamics provides important complementary information to recent rapid advances in CB_1_ structural elucidation through crystallography and cryo-EM. In particular MD enables the study of receptor dynamics without the C- or N-tail artifacts commonly necessary in these latter techniques. This study identified CB_1_ receptor components warranting more detailed investigation through ligand design and pharmacological assay, and lays a platform for further elucidating the molecular basis for signalling bias profiles of CB_1_ ligands. Further work to expand and refine this analysis to wider ligand libraries, and extension to drug discovery is ongoing.

The authors thank the University of Auckland for the award of a Doctoral Scholarship to DMJF, a Faculty of Science Research Development Fund award (3717371) to DPF and a contribution from the Computational Biology research theme and the NZ Marsden Fund, for support through grant 22-UOO-205 to DPF and MG. Part of this work was made possible by use of New Zealand eScience Infrastructure (NeSI) high performance computing facilities. New Zealand’s national facilities are provided by NeSI and funded jointly by NeSI’s collaborator institutions and through the Ministry of Business, Innovation & Employment’s Research Infrastructure programme. https://www.nesi.org.nz.

