## Supplementary Information for "Essential Dynamics of CB_1_ Receptor-Agonist Complexes: Implications for Signalling Bias" (from another version of this preprint: version 1, 2024-03-29)

**Figure S1**

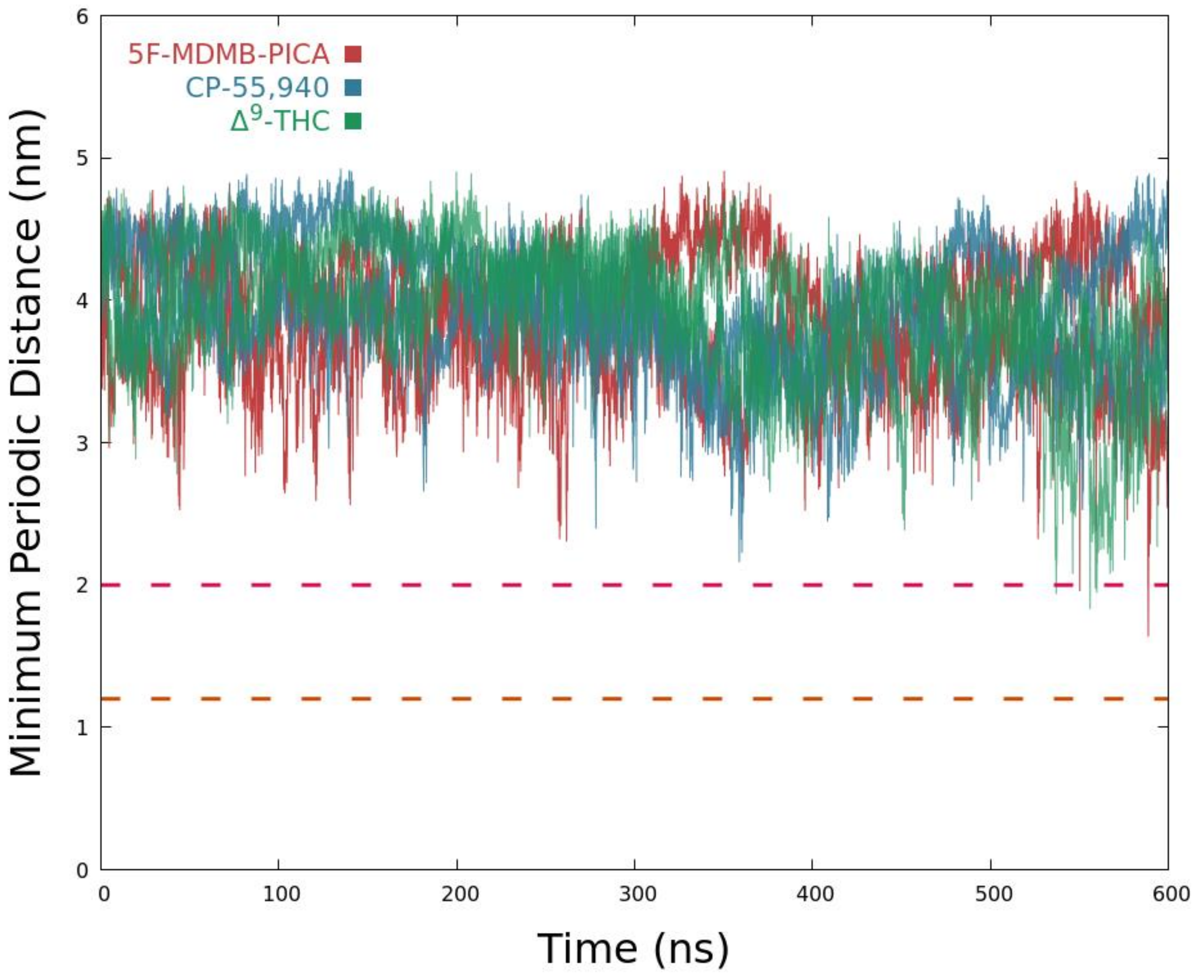


**Figure S1. Minimum periodic distance between receptor self-images over all 6 simulations.** The orange and pink dashed lines represent the 1.2 and 2.0 nm distances corresponding to the PME cutoff for nonbonded interactions, and the distance at which ordering effects of the protein on surrounding water/lipid become negligible. Apart from transient decreases below 2 nm in some trajectories near 550 ns, the receptor remains distant enough from its self-image to avoid becoming subject to unphysical forces which would invalidate the resulting trajectories.

**Figure S2**

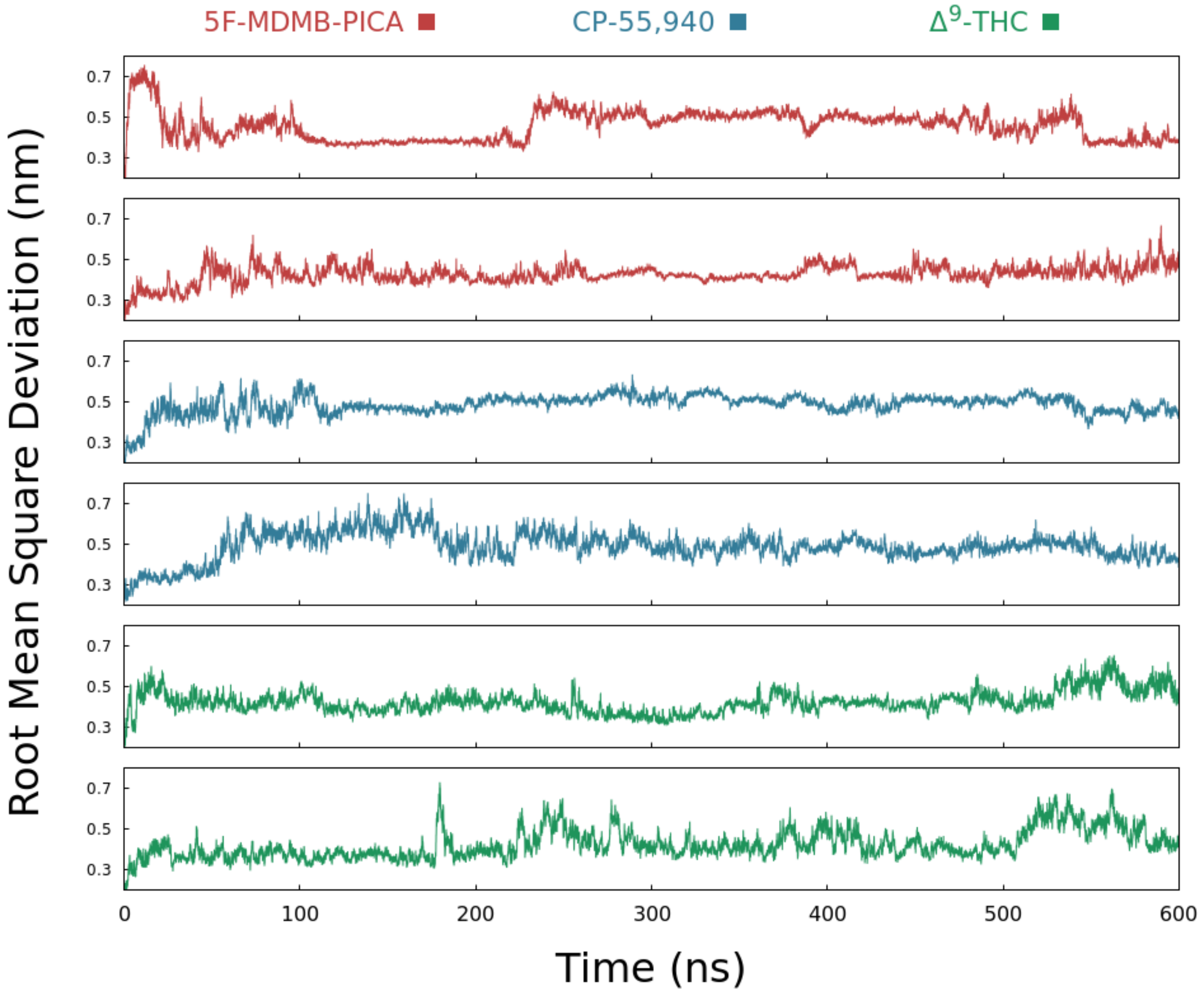


**Figure S2. Root Mean Square Deviation (RMSD) of the receptor compared to its initial conformation.** The RMSD values rapidly equilibrate to accommodate the presence of the ligand within the first tens of nanoseconds, after which each replicate reaches a more stable plateau. Sudden changes in the RMSD can be observed for some replicates, which indicate conformational changes in the receptor as part of its normal dynamics. Given enough simulation time, the ensemble of states should approach the ideal Boltzmann distribution for the receptor-membrane-ligand combination.

**Table S1.** Percentage contribution of the top ten principal components to the total variance in each group. Note: Trajectory frames of the regions were superimposed to themselves rather than the whole protein, to highlight intra-helical residue changes as opposed to gross protein motion (which is done separately, see Table ST2.)

| PC | TM1 | TM2 | TM3 | TM4 | TM5 | TM6 | TM7 | H8 | IC1 | IC2 |
|---|---|---|---|---|---|---|---|---|---|---|
| 1 | 10.2 | 18.7 | 20.7 | 18.9 | 11.6 | 22.8 | 13.8 | 14.7 | 43.7 | 27.6 |
| 2 | 7.3 | 10.0 | 12.4 | 8.1 | 8.6 | 8.5 | 12.8 | 12.0 | 19.1 | 16.9 |
| 3 | 5.6 | 5.3 | 5.0 | 6.1 | 7.5 | 5.5 | 5.9 | 8.6 | 7.9 | 11.2 |
| 4 | 4.5 | 4.0 | 4.4 | 3.9 | 5.6 | 4.6 | 4.1 | 7.6 | 6.3 | 8.6 |
| 5 | 3.9 | 3.8 | 4.0 | 3.8 | 4.4 | 4.3 | 3.9 | 7.2 | 3.2 | 3.5 |
| 6 | 3.7 | 3.5 | 2.8 | 3.4 | 3.8 | 3.5 | 3.4 | 6.8 | 2.2 | 3.3 |
| 7 | 3.2 | 3.3 | 2.7 | 2.7 | 3.0 | 3.0 | 3.2 | 4.0 | 1.7 | 2.7 |
| 8 | 3.0 | 2.9 | 2.2 | 2.4 | 2.6 | 2.6 | 2.8 | 3.4 | 1.3 | 2.0 |
| 9 | 2.8 | 2.3 | 2.0 | 2.1 | 2.4 | 2.3 | 2.8 | 3.0 | 1.2 | 1.6 |
| 10 | 2.5 | 2.2 | 2.0 | 2.0 | 1.7 | 2.0 | 2.5 | 2.3 | 1.1 | 1.4 |

**Table S2.** Percentage contribution of the top ten principal components to the total variance in each group. Note: Trajectory frames of the regions were superimposed to the whole protein, to highlight differences in sub-group positioning relative to the bundle.

| PC | TM1 | TM2 | TM3 | TM4 | TM5 | TM6 | TM7 | H8 | IC1 | IC2 |
|---|---|---|---|---|---|---|---|---|---|---|
| 1 | 36.7 | 26.2 | 21.3 | 41.0 | 25.0 | 32.9 | 34.4 | 40.2 | 37.6 | 39.4 |
| 2 | 12.2 | 11.7 | 18.6 | 14.1 | 18.1 | 18.0 | 12.6 | 14.4 | 18.8 | 23.9 |
| 3 | 8.3 | 7.7 | 10.0 | 8.3 | 8.2 | 6.9 | 8.8 | 10.0 | 15.6 | 8.0 |
| 4 | 7.2 | 7.0 | 7.6 | 5.1 | 5.1 | 5.6 | 6.2 | 5.6 | 6.4 | 4.8 |
| 5 | 3.3 | 5.5 | 5.1 | 3.4 | 4.2 | 5.5 | 5.1 | 4.3 | 4.5 | 4.0 |
| 6 | 2.3 | 3.6 | 3.6 | 2.5 | 3.6 | 3.5 | 3.3 | 3.0 | 2.6 | 3.2 |
| 7 | 1.9 | 3.1 | 2.3 | 2.0 | 2.8 | 2.8 | 2.3 | 2.3 | 2.4 | 2.7 |
| 8 | 1.7 | 2.5 | 2.2 | 1.6 | 2.2 | 1.8 | 1.8 | 2.2 | 1.9 | 1.8 |
| 9 | 1.5 | 1.9 | 1.9 | 1.3 | 1.9 | 1.5 | 1.5 | 1.9 | 1.5 | 1.5 |
| 10 | 1.3 | 1.7 | 1.5 | 1.2 | 1.7 | 1.3 | 1.3 | 1.6 | 1.0 | 1.2 |

### METHODS

#### Model preparation

The $CB_1$ cannabinoid receptor model used in this study was based on the active state cryo-EM $G_{i1}\alpha$-bound structure (PDB ID: 6N4B).[1] Missing residues in the N-terminus up to C98, and on the C-terminus up to C415 were filled in using MODELLER 9.21.[2] 100 models were generated and the highest scoring model was chosen for further refinement in Schrödinger Maestro 2020-2.[3] Truncated residues were fixed using Prime,[4,5] hydrogens were added, and disulfide bridges were formed. Protonation states were assigned using Epik,[6,7] the hydrogen-bond networks were optimised and the structure was subject to brief restrained refinement to settle clashes. Using Prime Loop, the N- and C-termini, ECL3, and ICL3 were modelled with a distance constraint on salt bridge residues K373-D176 on ECL3/TMH2.

After the initial 1 μs apo simulation, cluster analysis was performed based on receptor RMSD using the GROMOS clustering method as implemented in gmx cluster.[8] The representative structure of the major cluster in the latter 500 ns of the trajectory was extracted for use as the initial state of the ligand-bound simulations. Due to their structural similarity with the test ligands, the poses of the ligands in the cryo-EM template and a $CB_1$ active state crystal structure (PDB ID: 5XR8)[9] were used to superimpose the test ligands and insert them into the cluster structure. The resulting complexes were briefly minimised in Schrödinger Maestro to eliminate any clashes.

#### Small molecule parameters

The ParamChem server[10,11] was used to generate initial CGenFF[12] parameters. Atom typing was manually checked to ensure the most appropriate atom types were assigned for each ligand atom. Target data for parameter optimisation was generated in Gaussian 09RevD.01[13] using the MP2 method with the 6-31G* basis set for geometries, single-point energies, Hessian matrix, and relaxed dihedral scans. The HF method was used with the 6-31G* basis set for water-interaction scans. The Force Field Toolkit (FFTK) plugin for VMD 1.9.3 was used to optimise high-penalty partial charges and bonded parameters according to the published protocol.[14] Force field parameters were validated by minimising the ligands *in vacuo* with the test parameters and superimposing the resultant geometries with the MP2/6-31G* optimised geometries. In addition to comparing RMS deviations between the original and fitted parameters, individual bond lengths and angles were measured (With VMD 1.9.3)[15] and compared to those from the QM geometries to ensure deviations fell under 0.02 Å and 3° for bonds and valence angles, respectively. QM target data generation was carried out using Mahuika, the Cray CS400 HPC cluster provided by the New Zealand eScience Infrastructure (NeSI).

#### Molecular dynamics

The G protein chains were deleted and the receptor structure was submitted to CHARMM-GUI[16] Membrane Builder.[17,18] C-terminal residue C415 was palmitoylated to serve as a membrane anchor, and D338 was protonated to disfavour closure of the receptor-inactivating TMH3-6 salt bridge. The C-terminus was acetylated and N-terminus N-methylamidated. The PPM server was used to align the receptor to the membrane and a hetereogeneous bilayer composed of 282 lipids (POPC:SDPC:CHL1 in 7:7:6 ratio) was generated. The protein was inserted into the membrane and the system was solvated with TIP3 water, 0.15 M NaCl and neutralising ions, with a water thickness of 22.5 Å. The CHARMM36m parameters[19] were used with additional WYF parameters[20] to correct π-cation interactions. Hydrogen mass repartitioning[21] was enabled, allowing the use of a 4 fs time-step with LINCS restraints on hydrogens. The isothermal-isobaric ensemble (NPT) with periodic boundary conditions was used for the equilibration steps, while the production run was carried out in the canonical (NVT) ensemble, with the Nose-Hoover thermostat set to 310.15 K (37 °C) and Parrinello-Rahman barostat. The Verlet cutoff scheme was used with a 12 Å cutoff distance. All simulations and analysis were performed in GROMACS 2020.2.[22,23] After a 5000 step steepest descent minimisation, the system was initially equilibrated over 6 steps (3 x 125 ps at 1 fs time-step, and 3 x 500 ps at 2 fs time-step) with progressively relaxing restraints. A simulation of the receptor in the apo state was run for 1 μs in order to prepare the structure for the subsequent ligand-bound runs. Each of the three ligand-bound structures were simulated for 600 ns in duplicate (2 × 3 × 600 ns). All MD simulations and analysis were carried out on a local workstation with an nVidia RTX 2080 SUPER graphics card.

**Post-processing and analysis**

Trajectory processing and analysis was performed using GROMACS 2020.2. Trajectory visualisation and protein/ligand figure preparation was performed using PyMol 1.8.4[24] and VMD 1.9.3. Chemical structures were produced using ChemDraw 19.1. Graphs were produced using Gnuplot 5.2.[25] The first 100 ns from each ligand-bound 600 ns trajectory were discarded for analysis unless otherwise specified.

To align the protein structures for principal component analysis, all non-protein atoms were removed and a meta-trajectory of the 6 x 500 ns trajectories was produced with a rototranslational fit applied using the first frame as reference. The cartesian coordinates to be analysed were specified using custom index files produced by gmx make_ndx. Helix boundaries were defined by calculating the helicity per residue over the course of the simulations with gmx helix, and choosing a cutoff of contiguous helicity above 50%. The mass-weighted covariance matrix of the coordinates were calculated and diagonalised using gmx covar, and gmx anaeig was used to analyse the resultant eigenvectors. To highlight dynamic variance within each analysis group, the least squares fitting of the coordinates using gmx covar was performed using the initial structure of the analysis group as a reference, which we refer to as self-fit PCA. Conversely, to obtain data on the dynamics of the analysis group with respect to the protein bundle – i.e. to highlight helix displacement relative to the bundle – the least squares fit was performed using the whole protein structure as a reference, which we refer to as protein-fit PCA.

The RMS fluctuations per atom and extreme projections of the eigenvectors along the trajectory were used to determine the residue motions contributing to the major principal components, in order to identify residues which may be involved in ligand effects on signalling bias. The eigenvalues associated with each principal component were plotted and the point of maximum curvature was used as a threshold for retaining principal components for analysis, with some margin to ensure an even number of principal components were retained to facilitate viewing the data in 2 dimensions.

To gain a better understanding of the conformational changes described by the principal components, clusters of interest on the 2D plots were averaged and frames corresponding to the cluster centres were dumped using gmx trjconv, for visual inspection in PyMOL.

Ligand volume maps were calculated and visualised in VMD, utilising the VolMap plugin to calculate the average density of the ligand over the course of each simulation.

The GROMACS distance measurement tool gmx pairdist was used to obtain distance measurements between the ligands and each binding site residue, which were then classified as interacting or non-interacting for each frame of the trajectory based on whether the minimum distance was above or below 4 Å. Subsequently, the percentage of time the ligands were in contact with each residue over the simulation was calculated.
Ligand-receptor interaction energies were calculated by defining energy groups for each ligand and receptor subdomain and using the gmx rerun functionality to compute the energies based on the coordinates in the simulation trajectories. The interaction energy with each subdomain was normalised to the sum of that ligand's receptor interaction energy.

To obtain the data for rotamer analysis, the dihedral angles for residues of interest were calculated using PLUMED driver on the simulation trajectories.

Lastly, the gmx bundle analysis tool was used to calculate TMH7 kink angle and vertical displacement, for which the P394 residue of the NPxxY motif was defined as the kink-forming residue using a custom index group.

This work was carried out using the open-source, community-developed PLUMED library[26], version 2.7.[27]

**Acknowledgment**

The authors wish to acknowledge the use of New Zealand eScience Infrastructure (NeSI) high performance computing facilities, consulting support and/or training services as part of this research. New Zealand's national facilities are provided by NeSI and funded jointly by NeSI's collaborator institutions and through the Ministry of Business, Innovation & Employment's Research Infrastructure programme. URL https://www.nesi.org.nz.
